# Both behavior-manipulating and non-manipulating entomopathogenic fungi affect rhythmic gene expression in carpenter ant foragers upon infection

**DOI:** 10.1101/2023.01.19.524837

**Authors:** Biplabendu Das, Andreas Brachmann, Charissa de Bekker

## Abstract

**Background:** Behavioral plasticity in the nocturnal ant *Camponotus floridanus* is associated with changes in daily rhythms of core clock and clock-controlled genes in the brain. Plasticity in clock-controlled output, although adaptive, has been hypothesized to be a target for parasites that change host behavior in a timely manner to complete their life cycle. This study aims to explore this hypothesis by characterizing how the transcriptomic rhythms of the ant host change upon infection by a behavior manipulating parasite. We compared and contrasted the daily gene expression profile of uninfected *C. floridanus* ant heads to ants infected by a manipulating fungal parasite *Ophiocordyceps camponoti-floridani* and a non-manipulating fungus *Beauveria bassiana*, to test if changes to host clock and clock-controlled gene expression are specific to behavioral modifying diseases, or if such changes are a general hallmark of infectious diseases.

**Results:** The repertoire of genes oscillating every 24h in the ant heads showed almost three-fold reduction during *O. camponoti-floridani* infections, as compared to uninfected controls. Control-like nocturnal activity of 24h-rhythmic genes was maintained during *O. camponoti-floridani* infections, but not in *B. bassiana* infected ant heads. Half of all genes that showed 24h rhythms in the heads and brains of uninfected ants displayed highly synchronized changes in their rhythmic expression during both diseases, but in a species-specific manner. Network analyses revealed that both fungal parasites affected the same links between behavioral plasticity and clock output, albeit in a different manner.

**Conclusion:** Changes to clock-controlled transcriptomic rhythms of hosts might be a general hallmark of infectious diseases. However, the infection-associated changes to clock-controlled rhythms of the host are species-specific, and likely depends on the life history strategies used by the parasite.

## INTRODUCTION

Plasticity of the ant’s clock and clock-controlled rhythms seems to be tightly linked to its behavioral plasticity [1]. The ease with which ants display behavioral plasticity, although beneficial to the host, could make them susceptible to parasite hijacking. Parasites that modify host behavior in order to successfully grow and transmit are found in a wide variety of taxa. Field studies and laboratory experiments on different behavior modifying diseases indicate that manipulating parasites might be targeting the host’s biological clock to induce to some of the behavioral changes in its host. For example, flies infected with the fungus *Entomophthora* [2], caterpillars infected with baculoviruses [3], and ants infected with the fungus *Ophiocordyceps* [4] and the trematode *Dicrocoelium* [5] show altered, or even loss of, daily activity-rest rhythms. Furthermore, parasite-adaptive behaviors shown by infected hosts, such as summitting and attachment to substrate before death, seems to be highly synchronized to a specific time of day. Although the phenotypic evidence for a role of clocks in behavior modifying diseases seems to be emerging across different host-parasite systems, empirical evidence of the same on a molecular level is currently elusive.

An emerging model system to study parasitic manipulation of host behavior are the *Ophiocordyceps* fungi infecting ant hosts, and we focus primarily on one such parasite-host pair that is native to central Florida (USA): the carpenter ant *Camponotus floridanus* (host; *Cflo*) and their specialist manipulator *Ophiocordyceps camponoti-floridani* (parasite). As we layout in the General Introduction at the beginning of this dissertation, the *O. camponoti-floridani*-induced “manipulated” state in ants is associated with enhanced locomotory activity (or hyperactivity), increased wandering behavior, and severe convulsions. The characteristic manipulation phenotype is observed in the final stages of this disease: the infected ant shows a seemingly phototactic summiting behavior, and upon reaching a relatively elevated position, bites into a substrate with locked jaws, dying shortly after. The amount of light received at the final location of the manipulated ant seems to be important for *O. camponoti-floridani* fruiting body growth, whereas the elevated position has been hypothesized to be important for spore dispersal, and therefore, transmission of *O. camponoti-floridani* [6-8]. Taken together, the specific manipulation of ant behavior by *O. camponoti-floridani* provides a growth and transmission site that appears to be adaptive for the fungal parasite.

Moreover, the manipulated biting appears to be highly synchronized to a specific time of day across multiple *Ophiocordyceps*-ant species interactions, observed in field conditions as well as in controlled laboratory experiments [9-12]. This time-of-day specific manipulation of host behavior demonstrates that in the final stages of the disease, *O. camponoti-floridani*-infected ants do have some form of functional timekeeping machinery. It has been hypothesized that the fungal parasite might be driving at least some aspects of this timekeeping in the diseased ant [4]. Different lines of evidence suggest that *Ophiocordyceps* fungi could be hijacking the timekeeping machinery of the ant to induce timely manipulation of host behavior. First, like most fungi, *Ophiocordyceps* has a biological clock that is functionally similar to the endogenous oscillators found in animals and plants [13]. Second, several candidate fungal manipulation genes or effectors (e.g., small secreted proteins) seem to be under clock control in *Ophiocordyceps* [13]. And third, using a network-based meta-analysis, we have identified a possible molecular link between ant’s clock and behavioral plasticity that is likely targeted by the manipulating parasite to induce timely changes to the ant’s behavior and clock-controlled processes [4]. In light of the previous findings, we hypothesized that the manipulating parasite, *O. camponoti-floridani*, might be targeting a pre-existing regulatory link between the host ant’s clock and behavioral state in order to induce timely behavioral modifications [4]. One of the goals of the current study was to empirically test this apriori hypothesis by sampling *Ophiocordyceps*-infected ants over a 24h day to investigate what happens to the daily expression of their clock genes, clock-controlled genes and behavioral plasticity genes, and if the parasite does in fact affect the host’s pre-existing links between behavioral plasticity and clock.

Another goal of this study was to tease apart the changes that occur to the *C. floridanus’* clock and clock-controlled rhythms during infection by its manipulating, specialist parasite (*O. camponoti-floridani*) as compared to infection by a non-manipulating, generalist parasite (*Beauveria bassiana*). A synchronized timely change (increase/decrease) in the daily expression of behavioral plasticity genes can have multiple effects on the rhythmic properties of co-regulated clock-controlled genes, such as (1) loss or complete disruption of daily rhythms, (2) increase or decrease in the degree of synchronization of daily rhythms (e.g., most genes peaking at the same time of day), and (3) general increase or decrease in their average daily expression (differential gene expression). Such changes to host daily rhythms, however, could be a general effect of infectious diseases and not necessarily specific to manipulating parasites such as *Ophiocordyceps* (reviewed in [14]).

Previous work from our lab has characterized the effects these two infectious diseases have on host’s collective behavior, especially its ability to forage [15]. Trinh and colleagues demonstrated that in the early stages of *O. camponoti-floridani*-infection, prior to halfway through disease progression – the time point in disease at which only half of the infected ants survive – infected ants in a group perform extranidal (outside nest) visits in a rhythmic manner, similar to uninfected control groups [15]. However, in later stages of the disease, *O. camponoti-floridani*-infected ants show a loss of daily rhythms in foraging. It remains to be seen if this loss of rhythmic activity in group foraging is due to loss of individual locomotory rhythms in the infected hosts, or an inability of infected ants to detect or respond to social cues necessary for synchronizing their activity rhythms, or both. At least for *C. floridanus*, an absence of rhythms in locomotory behavior (e.g. arrhythmicity in nurse ants) does not necessarily mean an absence of functional timekeeping at the molecular level (several core clock and clock modulating genes oscillate every 8h in nurses) [1]. Unlike *O. camponoti-floridani*-infected ants, *B. bassiana-* infected ants do not show a loss of rhythmic foraging in the latter half of its disease. However, they do show a drastic shift in the daily timing of the foraging peak, shifting from a nocturnal peak during early disease, similar to uninfected controls, to a day-time peak around ZT6 (middle of the subjective day time) in the latter half of the disease [15]. Therefore, the halfway mark in both diseases seem to be a diverging point in their ultimate disease outcome, at least in terms of the parasite’s effect on host’s activity rhythms, which could be used to discern how daily gene expression is potentially affected by infection with both fungi.

In this study, we characterized the changes that occur to an infected forager ant’s clock and clock-controlled processes at the halfway mark in disease caused by *O. camponoti-floridani* as compared to *B. bassiana*. By combining time-course RNASeq and comparative network analyses, we aimed to (1) compare and contrast the changes in ant’s daily transcriptome caused by the two fungal parasites, (2) identify putative mechanisms via which *O. camponoti-floridani* might be inducing changes to the host’s clock and clock-controlled output, and (3) to test if manipulation of host’s daily gene expression is a hallmark of infectious diseases in general or specific to behavior manipulating parasites. The corresponding changes in *O. camponoti-floridani*’s daily transcriptome during infection, at the same disease stage, has been documented in a separate research article [16].

## MATERIALS AND METHODS

### Collecting Camponotus floridanus colony and experimental setup

We collected a large (several thousand workers), queen-absent colony of *C. floridanus* with abundant brood (eggs, larva, and pupa) from the University of Central Florida Arboretum on 19^th^ March 2021, from inside a decaying saw palmetto. Two days after collection, we split the source colony into two daughter colonies of approximately the same size and moved each into a clean container with talcum-coated walls. Until the start of the experiment, we kept the two daughter colonies under oscillating light-dark cycles (12h:12h) but constant temperature (25 ºC) and relative humidity (75% rH). Given that in natural conditions *Ophiocordyceps* is most likely to infect ants in the foraging caste (ones that perform most of the outside nest activities), we used only forager ants for fungal infections and as controls.

To identify the forager ants in a given daughter colony, we replicated the formicarium setup described in Das and de Bekker (2022) in which a foraging arena (container) was connected via a 2 m long plastic tube to another container always kept in dark [1]. The dark container imitated the dark nest chambers of an ant colony. Prior to starting the experiment, each individual daughter colony was moved into its own formicarium setup housed inside climate-controlled incubators. For the first phase of the experiment, the foraging arena of both daughter colonies were exposed to constant light conditions to incentivize ants to move their brood into the dark nest box. The constant light conditions in the foraging arena also aided in resetting the biological clocks of the ants and ensuring that the colony can later synchronize to the strict 12h:12h light-dark (LD) cycles they are exposed to. After 2-3 days of constant light conditions, the foraging arenas were exposed to 12h:12h LD cycles (lights were turned on at noon local time, and lights were turned off at midnight). Throughout the experiment, the colonies were kept at constant temperature (25 ºC) and relative humidity (75% rH). After three days of initial entrainment to the light-dark cycles, each daughter colony was sugar starved for at least 24h before we performed mark-and-recapture to identify foragers in the colony. Overnight sugar starvation helped with ensuring the colony was actively foraging during our days of mark-and-recapture in which we collected ants in the foraging arena during their active phase (night-time for our nocturnal ants), marked their gasters with a dot of silver paint (Testors) if collected the first time, or marked their thorax with a second dot of paint if collected for the second time. We performed all mark-and-recapture within two hours after lights were turned off (ZT12-ZT14) and two hours prior to lights turning on (ZT22-ZT24/ZT0). We then proceeded to setup smaller experimental colonies – one for controls and one for fungal infections – that contained either infected (with *O. camponoti-floridani* or *B. bassiana*) or uninfected (control) foragers. The experimental setups also ensured that the size of our experimental colonies was standardized. Each of our experimental boxes contained 150 marked forager ants that were either infected with fungi or served as uninfected controls. Additionally, we added 10 brood-tending nurse ants and some brood (10 larva and 10 pupa) to incentivize foraging during the experiment.

Due to logistical constraints, we could setup only two experimental colonies at a given time. For the first run, we constructed two experimental setups, one for infection trials with *B. bassiana* and another that served as control. Once *B. bassiana-*infected ants were sampled along with the controls for RNASeq, we setup two more experimental colonies – one for *O. camponoti-floridani* infection and the other served as control – to harvest infected ants at the same two-hour resolution used for *B. bassiana*-infected and control ants. For the *B. bassiana* infection run, we sampled the 300 foragers (150 for *B. bassiana-*infections and 150 for controls) from 445+ marked ants across the two daughter colonies that had visited the foraging arena at least once. For *O. camponoti-floridani* infection run, we sampled the 300 foragers from at least 378 marked ants.

*Camponotus floridanus* ants infected with *O. camponoti-floridani* can live up to four weeks before they show manipulated biting and ultimate death, whereas *B. bassiana* infected ants die within the first week post-infection. To standardize our comparison of the changes in ant daily gene expression during infection by *O. camponoti-floridani* as compared to *B. bassiana*, we sampled infected ants at halfway through disease progression (∼50% of infected ants in the colony had died). This is at the time during infection at which disruptions in daily activity rhythms have been observed for *O. camponoti-floridani*-infected ants [15]. For either infection run, we monitored disease progression by performing daily mortality checks and visualizing survival curves of the infected (and control) colonies. For *B. bassiana* infections, the disease reached halfway on Day 3 post-infections. Whereas *O. camponoti-floridani* infections reached the halfway mark on Day 11 post-infections (Additional File 3). As mentioned above, uninfected ants were collected from the control colony in parallel to sampling *B. bassiana-*infected ants. All the experimental runs were performed within a month (Apr 15^th^ to May 5^th^, 2022), under the same controlled climatic conditions (12h:12h LD, 25 ºC, and 75% rH). The climatic conditions of the foraging arena throughout the experiment were monitored using an environmental data logger (HOBO) (Additional File 2). The protocol used to monitor and quantify ant foraging rhythms are detailed in Das and de Bekker 2022 (see Methods section: *Colony activity monitoring*) [1].

### Fungal culturing and controlled infections

Fungal culturing and controlled infections of ant hosts were performed using the same methods and fungal strains as described in Trinh et al. (2021) [15]. For *O. camponoti-floridani* infections, we used the *O. camponoti-floridani* Arb2 strain, isolated from a manipulated *C. floridanus* ant cadaver previously collected from the wild [12]. For *B. bassiana* infections, we used the *B. bassiana* strain Bb0062 [17]. For both fungi, first we obtained fresh blastospores (i.e., yeast-like single cells) suspended in Grace’s Insect Medium (Gibco, Thermo Fisher Scientific) supplemented with 2.5% FBS (Gibco) using established protocols [12, 13, 18]. To infect ants, we injected (*O. camponoti-floridani*) or pricked (*B. bassiana*) ants with freshly harvested blastospores. As shown in Trinh et al. (2021), pricking ants with *B. bassiana* reliably caused infections while extending host’s survival time as compared to *B. bassiana*-injected ants [15]. For *O. camponoti-floridani* infections, we injected 0.5 µL of the blastospore solution (2×10^7^ cells/ml) into the ventral side of the thorax between its legs using 10 ml borosilicate capillary tubes (Fisher) fitted onto an aspirator (Drummond Scientific). Whereas for *B. bassiana* infections, we dipped the capillary tubes in the blastospore solution (2×10^7^ cells/ml) prior to pricking the ants on the same ventral side of its thorax. Control ants were neither injected nor pricked.

### Sample collection, RNA extraction, and RNA Sequencing

To obtain daily transcriptomes of host ant heads halfway through fungal infection, we sampled three infected (marked) foragers every 2h, over a 24h light-dark period, at 3 days post *B. bassiana-*infection and 11 days post *Ophiocordyceps-*infection. To obtain corresponding daily transcriptomes of hosts in an uninfected state, we sampled marked foragers from the control colony run alongside *B. bassiana-*infections, in parallel to sampling *B. bassiana*-infected ants, using the same sampling regime. Ants were collected in cryotubes, immediately flash frozen in liquid nitrogen, and stored at −80ºC until further use.

Prior to RNA extractions, for each treatment (control, *O. camponoti-floridani-*infected, and *B. bassiana-*infected ants), we pooled the three heads (antenna removed) harvested at each time point in a frozen cryotube containing two frozen steel ball bearings (5/32” type 2B, grade 300, Wheels Manufacturing). Next, we homogenized the pooled heads using a 1600 MiniG tissue homogenizer (SPEX) at 1300 rpm for 40 sec while keeping the samples frozen. Next, we isolated total RNA from the disrupted head tissues, followed by preparing cDNA libraries for RNASeq using the exact methods discussed in Das and de Bekker (2022) [1]. For each library preparation, we started with 500 ng of total RNA, extracted mRNA with poly-A magnetic beads (NEB), and converted this mRNA to 280-300 bp cDNA fragments using the Ultra II Directional Kit (NEB). Unique sequencing adapters were added to each cDNA library for multiplexing (NEB).

All thirty-six cDNA libraries were first sequenced as 50 bp single-end reads on an Illumina HiSeq1500. Given that fungal infected ant heads might contain mRNA from both ant and fungi, we re-sequenced the twenty-four mixed transcriptomes to obtain enough reads to be able to look at the daily transcriptomes of both ant host and fungal parasite (see [16]). The re-sequencing was performed as 75 bp paired-end reads on an Illumina NextSeq 1000. We combined the sequencing reads for the mixed transcriptomes by concatenating the raw reads from the two runs. Since we combined reads from single-end HiSeq run and paired-end NextSeq run, we used only the R1 reads from the latter.

### Processing RNASeq data and obtaining normalized gene expression

Prior to any analyses, we first removed sequencing adapters and low-quality reads from our RNASeq data using BBDuk (parameters: qtrim = rl, trimq = 10, hdist = 1, k = 23) [19]. Post-trimming, we used HISAT2 [20] to map transcripts to the relevant genomes. We used the publicly available genomes of *C. floridanus* (Cflo v7.5; [21]), *O. camponoti-floridani* (NCBI ID: 91520; [12]) and *B. bassiana* (ARSEF 2860; [22]). For all control *C. floridanus* samples, we mapped reads to the ant genome [23]. However, for fungal-infected *C. floridanus* samples, we first mapped all reads from the mixed transcriptomes to the respective fungal genomes to ensure that we only retained reads that are not of fungal origin, following which we mapped the remaining reads to the ant genome. Finally, we obtained normalized gene expression from the mapped reads as Fragments Per Kilobase of transcript per Million (FPKM) using Cuffdiff [24].

### Data analyses

We used the rhythmicity detection algorithm empirical JTK-Cycle (eJTK) [25, 26] to test for significant diurnal (24h) and ultradian (12h and 8h) rhythms in gene expression. Only genes that had diel expression values ≥1 FPKM for at least half of all sampled timepoints were tested for rhythmicity. For a set period length, a gene was considered to be significantly rhythmic if it had a Gamma p-value < 0.05.

As a proxy for a gene’s amplitude, we calculated its coefficient of variation throughout the 24h day. For a given gene, its coefficient of variation provides a normalized score of the variation observed in its expression around the mean. Therefore, using coefficient of variation as a proxy for amplitude allows us to compare the degree of daily fluctuations for any gene, not only the ones classified as rhythmic by a rhythmicity detection algorithm. For a given set of genes, we tested if the mean amplitude was significantly different between treatment groups using Kruskal-Wallis H test [27], and the post-hoc pairwise comparisons were done using Wilcoxon signed-rank test [28], using the “compare_means” function from the ggpubr package in R [29]. To cluster set of genes according to their daily expression, we used an agglomerative hierarchical clustering framework (method = complete linkage) using the ‘hclust’ function in the stats package for R.

To determine differentially expressed genes, we used the linear modelling framework proposed in LimoRhyde [30], but without an interaction between treatment and time. A gene was considered significantly differentially expressed if treatment was found to be a significant predictor (at 5% FDR) and the gene showed at least a two-fold change in mean diel expression between controls and *O. camponoti-floridani*-infected (or *B. bassiana*-infected) ants (abs(log_2_-fold-change) ≥ 1).

To perform functional enrichment analyses, we used an updated version of the custom enrichment function that performs hypergeometric test (previously used in [12] and [1]). The function “check_enrichment” is now publicly available for use via the timecourseRnaseq package on GitHub (https://github.com/biplabendu/timecourseRnaseq) [31]. For a given set of genes, we identified overrepresented Gene Ontology (GO) terms or PFAM domains using the check_enrichment function, and significance was inferred at 5% FDR. If not mentioned otherwise, for functional enrichment tests, we used all genes that were found to be “expressed” (≥ 1 FPKM expression for at least one sample) in the ant heads, for each treatment, as the background geneset. We only tested terms annotated for at least 5 protein coding genes. We used the GO and PFAM annotations [32] for the most recent *C. floridanus* genome (v 7.5) [23].

All data wrangling, statistical tests and graphical visualizations were performed in RStudio [33] using the R programming language v3.5.1 [34]. Heatmaps were generated using the pheatmap [35] and viridis [36] packages. Upset diagrams were used to visualize intersecting gene sets using the UpsetR package [37]. We used a Fisher’s exact test for identifying if two genesets showed significant overlap at 5% FDR. We visualized the results of multiple pairwise Fisher’s exact test using the GeneOverlap package [38]. All figures were generated using the ggplot2 package [39].

### Network analyses

To build the annotated gene co-expression network (GCN), we used the exact protocol detailed in de Bekker and Das (2022) [4] (Additional File). The clustering of genes into co-expressed modules was performed using functions from the WGCNA package [40-42], the gene-gene and module-module correlations were calculated using Kendall’s tau-b correlation [43], and the global connectivity patterns of the GCN was visualized using the igraph package [44]. The significance of pairwise overlaps were calculated using Fisher’s exact tests and visualized using the GeneOverlap package [38].

## RESULTS AND DISCUSSION

### General patterns of daily gene expression in ant heads

We defined genes to be “expressed” in *C. floridanus* ant heads if their mRNA levels were greater than 1 FPKM for at least one time point during the 24h sampling period. Of the 13,808 protein coding genes annotated in the *C. floridanus* genome [21], we found similar number of genes to be expressed in ant heads during infection as in healthy controls (73%, 75%, and 72% of *C. floridanus* genes were expressed in control, *O. camponoti-floridani*-infected, and *B. bassiana*-infected ants, respectively), and their pairwise overlaps were significant (Fisher’s exact test: odds ratio > 957 and p < 0.001). Of interest were 246 genes uniquely expressed in ant heads during *O. camponoti-floridani* infection (not expressed in controls or *B. bassiana*-infected ants), given that only 41 such uniquely expressed genes were found in *B. bassiana*-infected ants and 76 genes were uniquely expressed in control ants (Fig. 1A). The two distinct sets of uniquely expressed genes, in healthy controls and in *O. camponoti-floridani*-infected ants, were significantly enriched in the same olfactory processes; genes annotated with the Gene Ontology (GO) terms olfactory receptor activity, odorant binding, sensory perception on smell (Fig. 1B; Additional File 15A). Therefore, it seems that while several olfactory genes usually expressed in healthy ants are deactivated during *O. camponoti-floridani* infections, a distinctly different set of olfactory genes is activated simultaneously.

**Figure 1.**
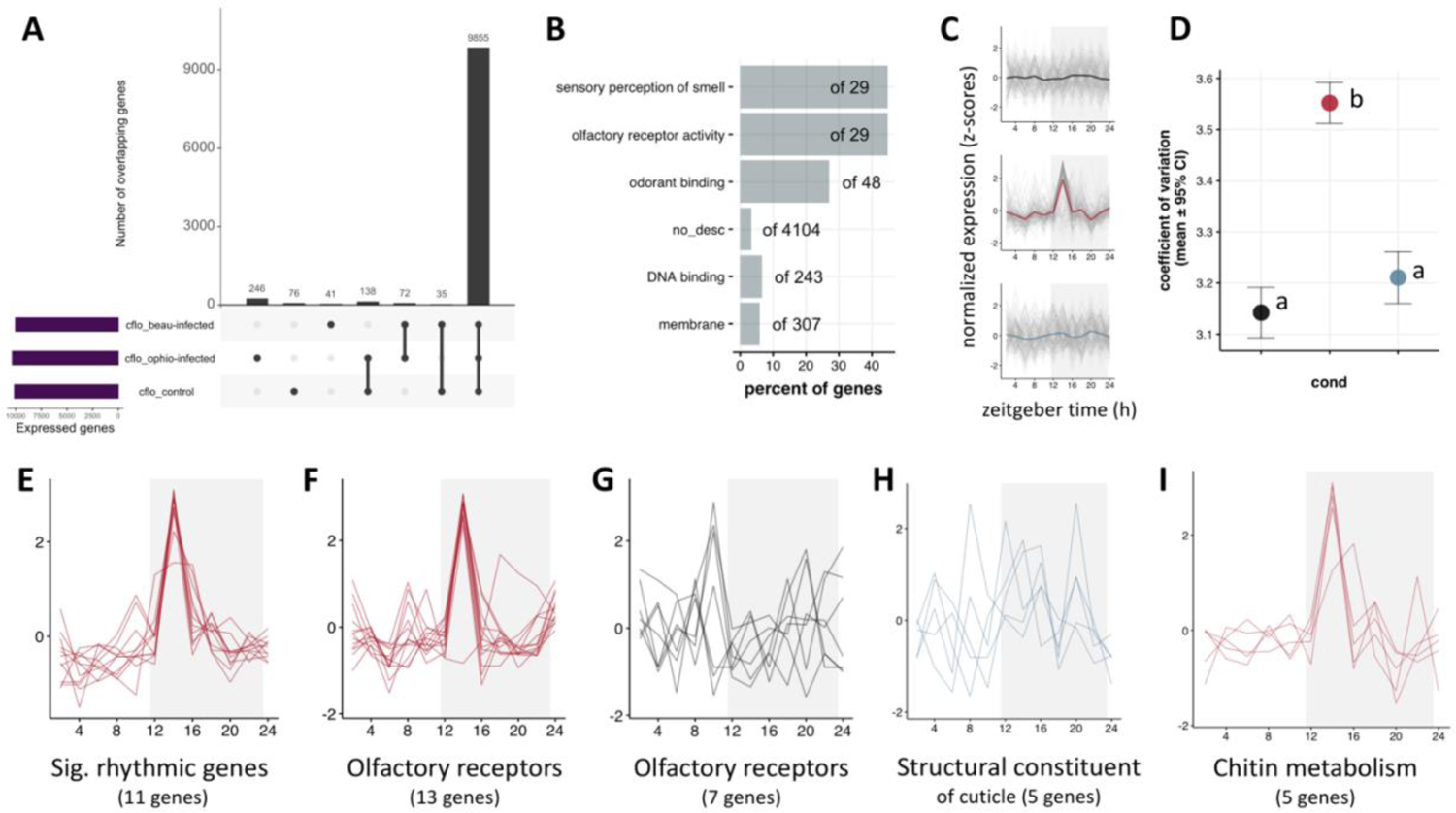
Uniquely expressed ant genes during O. camponoti-floridani infections are involved in olfaction. The panel (A) shows the number of genes expressed in ant heads during *B. bassiana* infections (cflo_beau-infected), *O. camponoti-floridani* infections (cflo_ophio-infected) and in uninfected control heads (cflo_control), as well as the number of genes intersecting between these three gene sets. Panel (B) shows the overrepresented GO terms in the 246 genes expressed uniquely during *O. camponoti-floridani* infections. The bars represent the percent of genes annotated with a given GO term that is found in the test gene set as compared to all such genes in the background gene set (in this case, all genes in the ant genome). Panel (C) shows the daily expression of the 246 genes uniquely expressed during *O. camponoti-floridani* infections, for each treatment group, as a “stacked zplot”. For a stacked zplot, the y-axis always represents the z-score normalized gene expression, and the x-axis represents zeitgeber time (in hour). Each grey line in the plot represents the expression of one gene, whereas the colored lines represent the median expression of all the genes used to build the stacked zplot. Throughout this manuscript, the color black is used to indicate data from uninfected foragers (control), red for *O. camponoti-floridani*-infected foragers, and blue for *B. bassiana*-infected foragers. Panel (D) shows the average (± 95% CI) amplitude of the same 246 genes shown in (C) for the three treatment groups. Colors have the same meaning. Different letters indicate significantly different amplitudes (p < 0.05). Additionally, stacked zplots are shown for (E) uniquely expressed genes during *O. camponoti-floridani* infections that are also classified as significantly rhythmic, (F) olfactory genes uniquely expressed in *O. camponoti-floridani*-infected ants, (G) olfactory genes uniquely expressed in uninfected controls, (H) genes annotated with the GO term structural constituent of cuticle that are expressed during *B. bassiana* infections but not in controls, and (I) genes annotated with the GO term chitin metabolism that are expressed during *O. camponoti-floridani* infections but not in controls.

In terms of daily fluctuations, most of the genes uniquely expressed in *O. camponoti-floridani*-infected ants showed a synchronized peak at ZT14 (two hours after lights are turned off) (Fig. 1C-D). Eleven of these uniquely expressed genes were also classified as significantly 24h-rhythmic in *O. camponoti-floridani*-infected ant heads including a putative serotonin receptor (*5-hydroxytryptamine receptor 3A*-like) and an anti-apoptotic protein coding gene (*death-associated inhibitor of apoptosis 1* like) (Fig. 1E). Although neither of the odorant receptor genes were classified as significantly rhythmic, the shapes of their daily expression profiles were almost identical to that of the significantly rhythmic genes (Fig. 1E-F). Neither our environmental data nor sequencing statistics indicate an abnormality for *O. camponoti-floridani*-infected ant samples collected at ZT14 (Additional File 3). Moreover, such a degree of time-of-day synchronization of daily expression was not observed for the uniquely expressed olfactory genes in controls, which were kept under the same conditions (Fig. 1G). Therefore, this synchronized timely peak observed for uniquely expressed genes in *O. camponoti-floridani*-infected ants is most likely a biological signal and not an experimental artefact.

In ants, like other insects, odorant receptors are known to regulate acceptance and avoidance behavior based on sensory perception of odor, e.g., response to pheromones in ant colonies. Additionally, colony odors can be a strong zeitgeber to entrain the clocks of social insects, even stronger than light-dark cycles [45]. Therefore, timed activation of the ant’s olfactory-mediated entrainment pathway can be a possible strategy for fungal parasites to regulate the phase of the ant’s clock and clock-controlled behavioral output. The 13 uniquely expressed olfactory genes in *O. camponoti-floridani*-infected ants contained the odorant receptors *OR10a, OR43a, OR85d*, an *OR2*-like gene, and two copies of *OR13*-like genes.

Whereas the seven olfactory genes uniquely expressed in controls, but not during *O. camponoti-floridani*-infection, contained copies of *OR13a*-like, *OR82a*-like, *OR82a, OR4*-like, and *OR4* genes. Although not much is known about the function of these odorant genes in social insects, a previous study in the fruit fly *D. melanogaster* has shown that differences in expression levels of *OR10a* and *OR43a* correlate with differences in fly response (attraction v. repulsion) to an aversive odor (benzaldehyde) [46]. Therefore, a synchronous daily peak expression of olfactory genes in *O. camponoti-floridani*-infected ant heads might correlate with a drastically reduced or heightened sensitivity of infected ants to certain colony odors in a time-of-day specific manner. Taken together, the data suggests that in *O. camponoti-floridani*-infected forager ants, the olfactory pathway might be playing a role in setting the phase of the host ant’s clock and its rhythmic output. In the next section, we have attempted to answer this question, at least partially, by testing (1) if a significant number of 24h-rhythmic genes in *O. camponoti-floridani*-infected ants show a daily peak (or trough) in expression at ZT14, (2) how the distribution of the phases (peak time-of-day) of significantly rhythmic genes compared across the three treatment groups, and (3) if the functions of rhythmic genes that show a waveform similar to the uniquely expressed olfactory genes during *O. camponoti-floridani*-infection are known to be under clock-control in ants.

### Daily rhythms in gene expression: forager heads vs. dissected brains

We identified rhythmic gene expression in ant heads using the non-parametric algorithm empirical JTK Cycle (eJTK) [25, 26]. Of the approximately ten thousand genes expressed in *C. floridanus* forager heads, 8.4% (852 genes) showed significant 24h-rhythmic expression in uninfected controls (Additional File 16A). At halfway through *O. camponoti-floridani* infection, however, only 2.9% (294 genes) of all expressed genes showed significant 24h-rhythms, whereas 6.7% (673 genes) showed significant diurnal rhythms at halfway through *B. bassiana* infection (Additional File 16B-C).

To validate our data, we compared the 24h-rhythmic genes identified in uninfected forager heads (this study) to those previously identified in forager brains [1]. We expected to see a reduced yet significant overlap in rhythmic genes between the brain and the head due to any of the following scenarios. First, even if a gene is rhythmic in every cell in the ant head, it might be oscillating with different phases in different tissues. Therefore, if there is enough tissue-specific variation in the phase of a rhythmic gene, we might not see a synchronous daily rhythm for the gene’s expression by sampling the entire head. Second, certain genes might be only rhythmic in the brain, and not in the surrounding muscle or gland tissues, which could dampen the amplitude of oscillating genes to the level where they are not readily detected. However, in the case that we do see rhythmic expression for a given gene in both ant brains and heads, we can be certain that it oscillates in both and is suggestive of a relatively high synchronization of its daily expression across different tissues inside ant heads.

A total of 334 genes were found to be 24h-rhythmic in both ant brains and whole heads. This overlap was significant (Fisher’s exact test: odds-ratio = 1.27, p < 0.001). However, no significant overlaps were found for genes oscillating every 12h or 8h in ant heads and ant brains. The reduced overlap for ultradian rhythms at the two different scales (brain tissue and whole heads) might be caused by the same reasons we discussed above. Indeed, gaining ultradian rhythms in whole ant heads as compared to brain tissue alone is difficult to interpret. They might be due to 24h-rhythmic genes that cycle with significantly different phases in different tissues, therefore, seeming ultradian when their expression is detected together. As such, we focused our attention on characterizing the changes that occur to the 24h-rhythmic gene expression of ants during infectious disease as compared to controls and interchangeably use the terms “rhythmic” and “24h-rhythmic” throughout.

### Changes in ant daily gene expression during infection: Differentially Rhythmic Genes

We wanted to explore the changes that occur to the rhythmic properties of genes that oscillate strongly enough to be rhythmic in both, brains [1, 4] and whole heads (this study) of foragers. The daily transcriptomes of forager heads and brains were sampled using the same experimental design, sampling resolution, and entrainment conditions, but the experiments were conducted a year apart using two different ant colonies. In the former study, comparison of the daily transcriptomes of forager and nurse ant brains revealed a set of behavioral-caste-associated differentially rhythmic genes (DRGs): 281 genes including the core clock *Period* and several clock modulators such as *Sgg, Dbt, Nemo, Pp1, Pp1b* that showed highly synchronized switch from oscillating every 24h in forager brains to oscillating every 8h in nurses (see Figure 3 in [1]). Therefore, it appears that plasticity of ant’s rhythmic (behavioral) state, and associated caste identity (task specialization) in the colony, is linked to synchronized changes in the daily expression of these 281 rhythmic genes. In this section, we wanted to test if the genes oscillating in both brains and heads show any signs of disease-associated differential rhythmicity (i.e., synchronized changes in daily gene expression due to disease), and if so, do these disease-associated DRGs show an overlap with the behavioral-caste-associated DRGs we have identified previously.

Among the 334 genes that show robust 24h oscillations in both ant brains and heads, hierarchical clustering revealed a set of 166 genes that show highly synchronized changes in their daily expression (Cluster-1, Fig. 2A-B). One could argue that we could have explored Cluster-2 as well, given that it also seems to show synchronized changes in daily expression. However, we chose to explore the genes in Cluster-1 because this is the largest cluster for which we found evidence of synchronized changes. The size of the cluster matters since our eventual goal was to test the possibility that disease-associated DRGs, if any, significantly overlapped with the caste-associated DRGs or not. As the cluster size would decrease, the chances of a false negative for a Fisher’s exact test would increase. That is why we decided to explore the 166 genes in Cluster-1. These 166 genes show a dawn peak in the forager brains (ZT22-24), whereas in forager heads we found a dusk peak (ZT12-ZT16) (Fig. 2A-B). The phase difference for these 166 genes between forager heads and brains could be, in part, due to the presence of tissue-specific variation in the phase of these genes throughout the entire head. Although these genes show a dawn phase in the brains, at the scale of the head, the various tissue-specific phases seem to integrate and produce a bell-shaped curve peaking around dawn. Regardless of the reasons for which these 166 genes show a phase shift between forager brains and heads, the fact that they can show a highly synchronized change in their daily rhythm is what motivated us to look at what happens to these genes during disease. We found a highly synchronized change in the shape of daily expression for these 166 genes during disease, as compared to controls (Fig. 2B).

**Figure 2:**
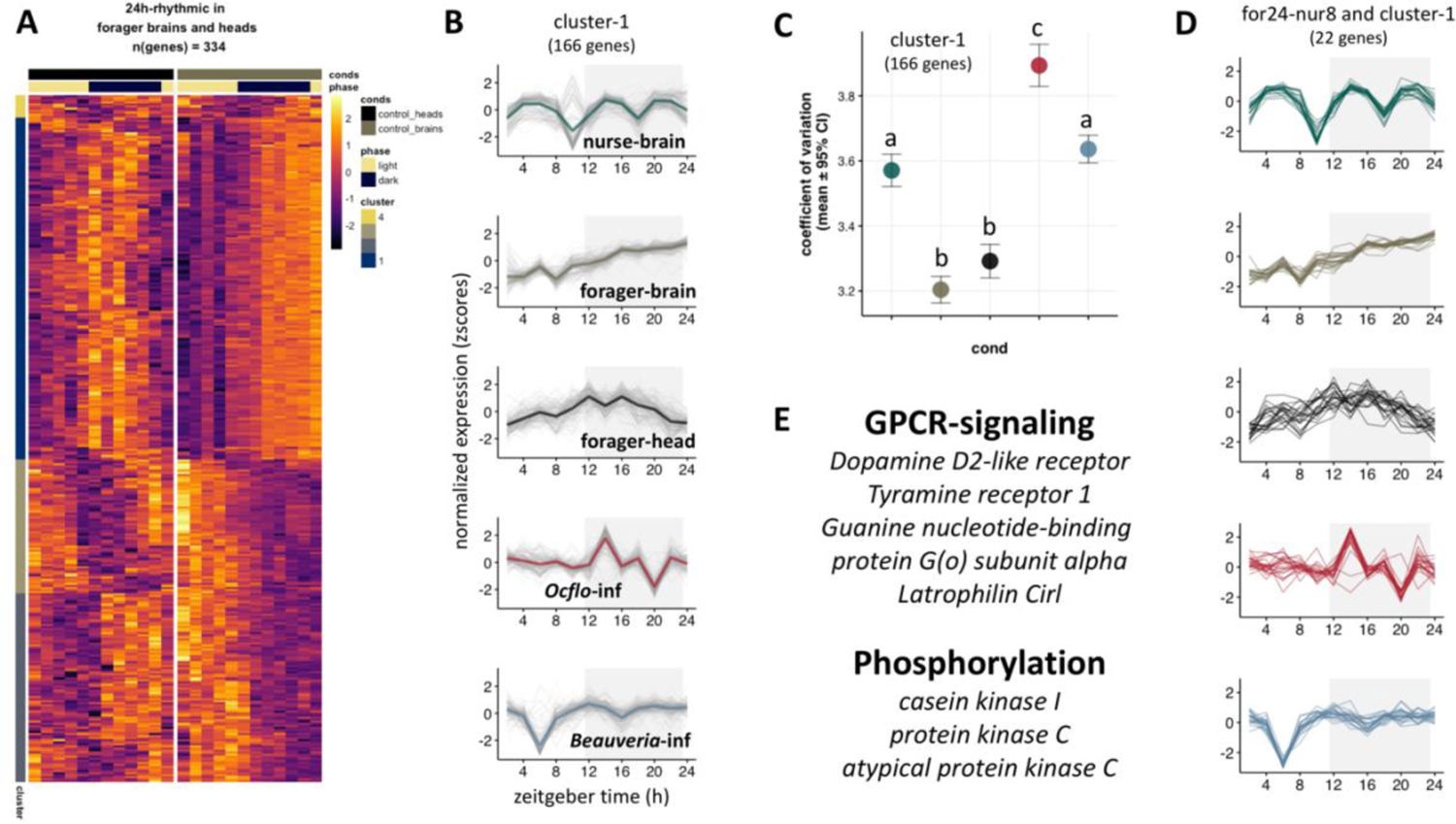
Genes that show 24h-rhythmicity in forager heads and brains show a synchronized change in daily expression during fungal infections, in a species-specific manner. The heatmap in panel (A) shows the daily expression (z-score) patterns of 334 genes identified as significantly oscillating every 24h in both forager heads and brains. Each row represents a single gene and each column represents the Zeitgeber Time (ZT) at which the sample was collected, shown in chronological order from left to right (from ZT2 to ZT24, every 2 h). These 334 genes were hierarchically clustered into four clusters, and the cluster identity of each gene is indicated in the cluster annotation column on the left of the heatmap. Panel (B) shows the stacked zplots for all 166 genes in Cluster-1 of the heatmap in (A). The data for nurse brains and forager brains were obtained from Das and de Bekker (2021). We have used the colors green and grey, respectively, to indicate the data for nurse and forager brains throughout the manuscript. In panel (C) we have plotted the mean (± 95% CI) amplitude of these 166 genes in Cluster-1. Different letters indicate significantly different amplitudes (p < 0.05). Panel (D) shows the stacked zplots for the 22 genes that were overlapping between the 166 genes in Cluster-1 and the 281 differentially rhythmic genes (for24-nur8) identified in Das and de Bekker (2021). Finally, in panel (E) we have highlighted two GO terms that were overrepresented in the 166 genes in Cluster-1, and some of the genes that contributed to the respective enrichment.

Furthermore, when we compared these 166 genes to the 281 caste-associated DRGs (for24-nur8), we found a significant overlap (Fisher’s exact test; odds ratio = 5.7, p < 0.001). Taken together, these two finding suggests that the core set of caste-associated DRGs in *C. floridanus*, which usually functions as molecular links between behavioral plasticity and plasticity of the clock, also shows synchronized changes in its daily expression during disease. Furthermore, the synchronized change at halfway through disease shows parasite-specific difference that might have a biological meaning. For example, the expression of these 166 differentially rhythmic genes peak around ZT14 in both uninfected forager heads and foragers infected with *O. camponoti-floridani*, but switches from smooth fluctuations in controls to a synchronized sharp peak (activation) in *O. camponoti-floridani*-infected ants (Fig. 2B). The same genes, however, do not show a clear peak around ZT14 in *B. bassiana*-infected ants. Instead, their daily expression shows a sharp dip (deactivation) at ZT6 with no apparent daily fluctuations during the rest of the 24h day (Fig. 2B).

This set of 166 disease-associated DRGs (brain-head-rhy24-cluster1) showed a significant overrepresentation for genes involved in protein phosphorylation and GPCR signaling pathway (Fig. 2E). The phosphorylation genes included *casein kinase 1*, both conventional and atypical *protein kinase C*, serine/threonine-protein kinase *SIK3, Calcium/calmodulin-dependent protein kinase II* (*CaMKII*), and *mitogen-activated protein kinase 1* (*MAPK1*; synonymous to *extracellular signal-regulated kinase 2* or *ERK2*). Whereas genes involved in GPCR signaling included a *dopamine D2-like receptor* and *tyramine receptor 1* (*TAR1*). In summary, key regulators of signal transduction (PKCs), insulin signaling pathway (serine/threonine protein kinases), synaptic plasticity (CAMKs), and social behavior (ERK2 and dopamine/tyramine receptors) show synchronized change in their daily expression during infectious disease, albeit in a parasite-specific manner.

We additionally asked how changes to the degree of daily fluctuation of these 166 disease-associated DRGs might correlate with the parasite-specific disease outcome we see in infected ants. To obtain a standardized metric for the degree of daily fluctuation observed for a gene, we calculated its coefficient of variation throughout the 24h day, which we refer to as the amplitude. For a given gene, its coefficient of variation provides a normalized score of the variation observed in its expression around the mean. Therefore, using coefficient of variation as a proxy for amplitude allows us to compare the degree of daily fluctuations for any gene, not only the ones classified as rhythmic by a rhythmicity detection algorithm. It appears that in uninfected conditions, nurse ants show a significantly higher amplitude of daily expression in these DRGs as compared to foragers (Wilcoxon signed-rank test: p-value < 0.001; Fig. 2C). As such, significant differences in the amplitude of these DRGs, in addition to differences in periodicity, seem to correlate with different behavioral state (nurse v. foragers). Given that the DRGs show a synchronized peak (or a trough) in infected foragers, we expected to see an increase in the amplitude of daily fluctuations in both diseased states as compared to uninfected foragers, which is what we found (Wilcoxon signed-rank tests: p-values < 0.001; Fig. 2C).

Furthermore, the disease-associated DRGs showed a significantly higher amplitude in *O. camponoti-floridani*-infected ant heads as compared to *B. bassiana*-infected ones as well as nurse brains (Wilcoxon signed-rank tests: p-values < 0.001; Fig. 2C). The amplitude of these genes in *B. bassiana*-infected ants were not significantly different than that of nurses (Wilcoxon signed-rank tests: p-values = 0.056; Fig. 2C). Therefore, it seems that the disease-associated changes to the behavioral state of the ants are associated with not only changes to the shape, but also the amplitude, of these 166 genes. Taken together, this suggests that in ants infected with *O. camponoti-floridani*, the peak time of daily expression of several essential clock components (*CK1* and *PKCs*) and behavioral regulators (dopamine and tyramine receptors) still mimics that of controls, albeit with a significantly higher degree to daily fluctuation. During *B. bassiana* infections, such an expression peak around ZT14 is lost, and rather a sharp dip at ZT6 is observed. As we discussed previously, although *B. bassiana*-infected ants seem to maintain robust foraging rhythms in the second half of the disease, the peak time of their daily foraging activity shifts from a night-time peak (usually between ZT12-16) to the middle of the day, at ZT6. It is unclear, but the possibility remains that the sharp deactivation of these 166 disease-associated DRGs at ZT6 might underlie the altered foraging peak seen in *B. bassiana*-infected ants. It also remains to be seen what drives this synchronized activation and deactivation, respectively, of these 166 disease associated DRGs during *O. camponoti-floridani* and *B. bassiana* infections.

### Ant gene expression during infection: Temporal division of clock-controlled processes

Of the 24h-rhythmic genes found in *O. camponoti-floridani*-infected ant heads, a majority showed a daily peak at ZT14 (two hours post lights-off), which was close to the phase of rhythmic genes in control heads (around ZT16) (Fig. 3A-B). During *B. bassiana*-infection, however, most 24h-oscillating genes in the infected ant’s head showed a peak expression at ZT6 (middle of the light phase) (Fig. 3C). The shift in the average phase of rhythmic genes in ant heads during infection with *O. camponoti-floridani* (Watson test statistic = 1.4, p < 0.001) and infection with *B. bassiana* (Watson test statistic = 7.7, p < 0.001), as compared to controls, were both significant. The phase shift in host’s clock-controlled genes was significantly higher during *B. bassiana* infection as compared to *O. camponoti-floridani* infection (Watson test statistic = 3.7, p < 0.001). This finding is consistent with the difference in clock-controlled activity rhythms that Trinh and colleagues found for ants infected with *O. camponoti-floridani* versus *B. bassiana. O. camponoti-floridani*-infected ants seem to maintain daily rhythms in foraging – a clock-controlled process – until the halfway mark in disease but lose such rhythmic activity in the second half of the disease. Given that the incubation period of *O. camponoti-floridani* inside its host is relatively long (around three to four weeks), and that we have sampled *O. camponoti-floridani*-infected ants exactly at the halfway mark in disease progression, we have most likely sampled infected ants that are in the process of transitioning from a seemingly rhythmic phenotype to an arrhythmic one. In other words, several clock-controlled processes in the host might still be oscillating in a rhythmic manner with a phase similar to uninfected controls. The latter, at least, seems likely since we found a similar phase of rhythmic gene expression in both uninfected controls and *O. camponoti-floridani*-infected ants. In comparison, the disease progresses relatively faster in *B. bassiana*-infected ants; infected ants succumb to the fungal infection within a week. Therefore, our sampled ants at the halfway mark might already give us a glimpse of what is about to happen to the infected ants in the latter half of the disease. The fact that we find that majority of the clock-controlled genes in *B. bassiana*-infected ants peak at ZT6, the same time of day at which their daily activity peaks during the latter half of the disease, suggests that by the halfway mark in its disease, *B. bassiana* likely causes a drastic phase shift in most host processes under clock control, including activity rhythms. The vastly different rate of disease progression of these two infectious diseases in ants, therefore, seems to be associated with the different degrees to which the host’s temporal division of clock-controlled processes is affected at the halfway mark in the disease. Future studies on other infectious diseases in ants, with varying rates of disease progression would be necessary to confirm our speculation.

**Figure 3:**
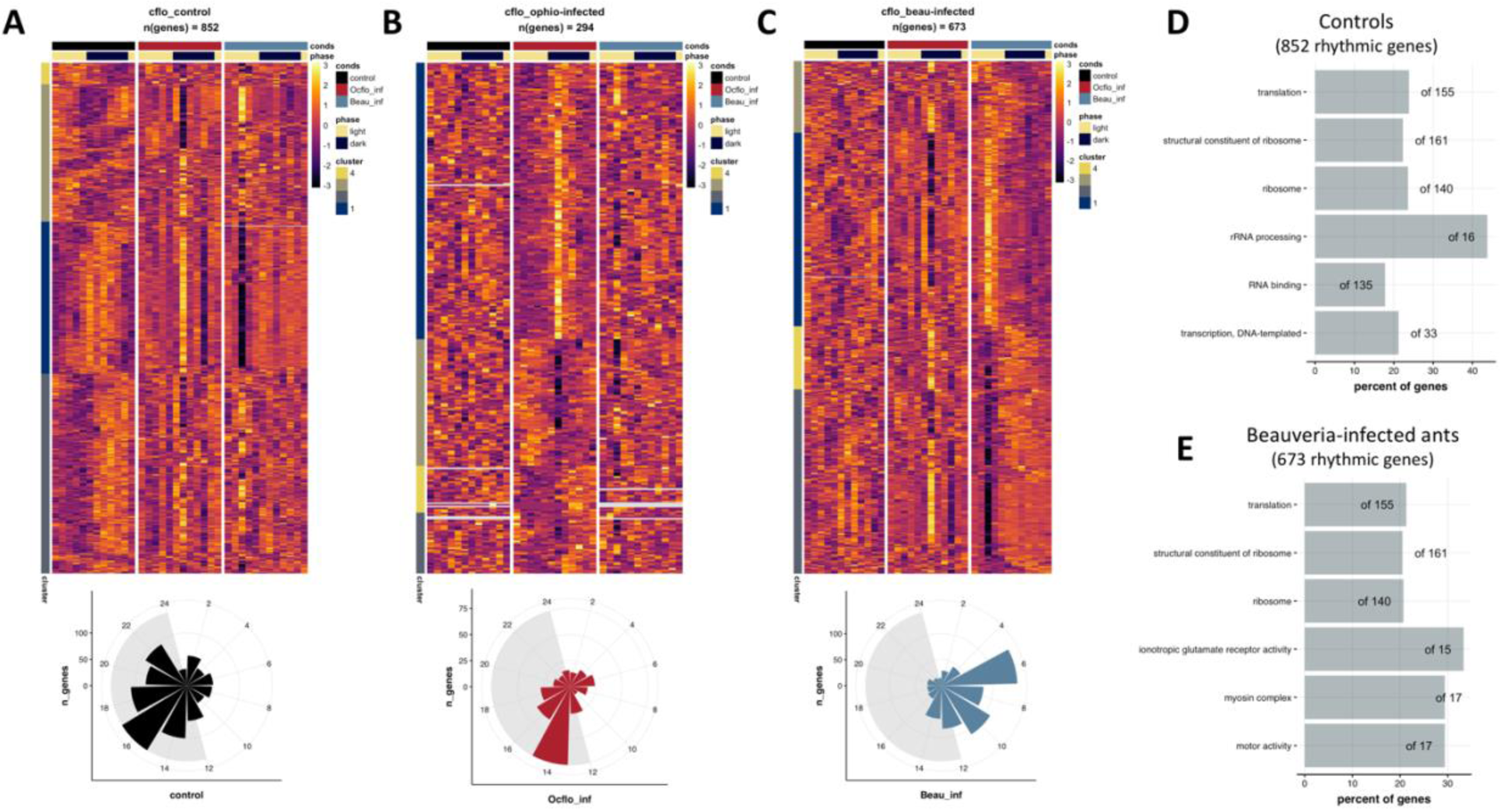
Clock-controlled biological processes are disrupted during O. camponoti-floridani infections but keep oscillating during B. bassiana infections, albeit with a drastic phase shift. The heatmaps show the daily expression profiles of genes identified as 24h-rhythmic in (A) uninfected controls, (B) *O. camponoti-floridani*-infected ants, and (C) *B. bassiana*-infected ants, whereas the phase plot at the bottom shows the distribution of the phases (time-of-day of peak expression) of the identified rhythmic genes. The corresponding daily expression of the same genes for the other two conditions are also shown as a reference. For example, (B) shows the daily expression pattern of the 294 genes that were identified as significantly rhythmic in *O. camponoti-floridani*-infected ant heads. For comparison, the daily expression of these 294 genes in uninfected controls and *B. bassiana*-infected ants are also shown. At the bottom of the heatmap, the distribution of the phases for these 294 rhythmic genes are shown. Panels (D) and (E), respectively, show the GO terms overrepresented in the genes identified as 24h-rhythmic in controls and *B. bassiana*-infected ants. No enriched GO terms were found for genes rhythmic in *O. camponoti-floridani*-infected ants. Uninfected control = cflo_control, *O. camponoti-floridani*-infected ants = cflo_ophio-infected, and *B. bassiana*-infected ants = cflo_beau-infected.

Next, we wanted to find out if the identity of the genes that are under clock control in uninfected foragers, and the processes they regulate, changed drastically at the halfway mark in either infectious disease. We expected to see a drastic change in the clock-controlled rhythms for *O. camponoti-floridani*-infected ants since both previous work [15] (Fig. 9) and activity data collected during this study (Additional File) showed a loss of foraging rhythms at this disease stage. However, for *B. bassiana*-infected ants, we expected to see less drastic changes to the repertoire of rhythmic genes, at least for ones underlying locomotion, since *B. bassiana*-infected ants tend to maintain rhythmic activity even in the latter half of the disease, although with a large phase shift. Using pairwise Fisher’s exact tests, we found that the set of 24h-oscillating genes in control forager heads did not significantly overlap with those identified as 24h-rhythmic in *B. bassiana*-infected or *O. camponoti-floridani*-infected ants. Although not significantly overlapping, the rhythmic genes identified in uninfected controls and *B. bassiana*-infected ants showed a higher overlap (63 genes) than those between controls and *O. camponoti-floridani*-infected ants (12 genes). Only three genes (*centromere-associated protein E, neither activation not afterpotential protein C* (*ninaC*), and *selenoprotein P-like*) were classified as rhythmic in all three conditions, and all three displayed a similar night-time peak in daily expression. Indeed, majority of the genes (780 genes in controls, 591 genes in *B. bassiana*-infected ants, and 263 genes in *O. camponoti-floridani*-infected ants) were classified as rhythmic in one treatment group but not the other.

Even though the identity of the rhythmic genes seems to change drastically in both disease conditions, their functions might be comparable. However, we did not find any enriched GO terms in among the 294 rhythmic genes in *O. camponoti-floridani*-infected ants. In contrast, the rhythmic genes in controls and during *B. bassiana*-infections did contain overrepresented gene functions for similar translation-related processes (Fig. 3D-E). This overlap was found to be significant (Fisher’s exact test: odds-ratio = 118, p-value < 0.001) (Fig. 2D-E). Therefore, it seems that although the clock-controlled genes shift from a night-time peak activity in uninfected controls to a mid-day peak (at ZT6) during *B. bassiana* infection, the rhythmic genes in *B. bassiana*-infected ants show a significant functional overlap with those oscillating in controls. The disease phenotype of *B. bassiana*-infected ants is likely due to the drastic effects *B. bassiana* has on the host’s temporal division of clock-controlled processes.

In addition to translation-related processes, the rhythmic genes in *B. bassiana*-infected ants were overrepresented in motor activity. The five genes involved in motor activity showed a synchronized daily expression in *B. bassiana*-infected ants with a sharp dip observed around ZT6, similar to the disease-associated DRGs discussed previously (Fig. 2B). These motor related genes included a *myosin heavy chain*, a *dilute class unconventional myosin*, a *paramyosin*, and *ninaC*. In *Drosophila*, these genes seem to be involved in muscle contraction [47], transport of pigment granules in photoceptors [48], general muscle integrity and function [49, 50], and photoreceptor cell function [51, 52]. Given that these genes seem to link photoreception to muscle function, it remains to be seen if the sharp trough in the daily expression of these motor-related genes at ZT6 correlates with an increased motor activity observed for *B. bassiana*-infected ants.

In summary, it appears that infection by both, manipulating and non-manipulating, fungal parasites can largely disrupt the identity of the clock-controlled genes in the host’s head. Although the identity of the 24h oscillating genes inside ant heads were distinct during infection with the biotrophic, specialist manipulator *O. camponoti-floridani*, the bulk of the rhythmic genes still showed a night-time peak of daily expression similar to uninfected controls. In comparison, infection by the necrotrophic, non-manipulating fungal pathogen *B. bassiana* caused relatively less drastic changes to the ant’s repertoire of rhythmic genes but induced a nocturnal to diurnal phase shift in the ant’s clock-controlled processes, consistent with its observed disease phenotype [15]. This large phase shift in the ant’s rhythmic output might be a hallmark of ants succumbing to *B. bassiana* infection, and in part, could be responsible for the relatively fast disease progression observed (*B. bassiana* kills its ant host within a week and does not require its host to be alive for extended periods to complete its lifecycle). In comparison, ants infected by *O. camponoti-floridani* need to survive for more than two to three weeks. This relatively long incubation time of *O. camponoti-floridani* is likely achieved by its host being able to maintain a control-like temporal structure of clock-controlled processes during disease, at least till the halfway mark. If *O. camponoti-floridani* aids in this process, by synchronizing to and maintaining its host’s rhythms, or not remains to be seen.

### Ant gene expression during infection: Differentially expressed genes

To discuss genes and biological processes that are drastically altered in ant heads during infection by fungal pathogens and characterize the differences between infection by *O. camponoti-floridani* and *B. bassiana*, we identified the genes that were significantly differentially expressed in ant heads between control and diseased conditions (fold change ≥ 2, q-value < 0.05). Of the approximately ten thousand genes expressed in ant heads, 143 were upregulated, and 81 were downregulated at halfway through *O. camponoti-floridani* infection as compared to controls (Fig. 4A). At a similar disease stage in *B. bassiana*-infected ants, the opposite trend was found: 80 ant genes were upregulated, and 139 were downregulated as compared to controls (Fig. 4B). We found significant overlap between ant genes downregulated during *O. camponoti-floridani* infection and ones upregulated during *B. bassiana* infection (*Ocflo*-DOWN-Beau-UP: 63 genes; odds ratio = 106, q-value < 0.001) (Fig. 4D). A smaller, yet significant, overlap was found between genes upregulated during *O. camponoti-floridani* infection and downregulated during *B. bassiana* infection (*Ocflo*-UP-Beau-DOWN: 18 genes; odds ratio = 47, q-value < 0.001) (Fig. 13D). The findings suggest that there exist two core sets of ant genes both of which show drastic changes in their expression during fungal infections, but in a parasite-specific manner. Whereas infection by *O. camponoti-floridani* causes upregulation (or downregulation) in a set of host’s gene, infection by *B. bassiana* causes downregulation (or upregulation) in the same set of host genes.

**Figure 4:**
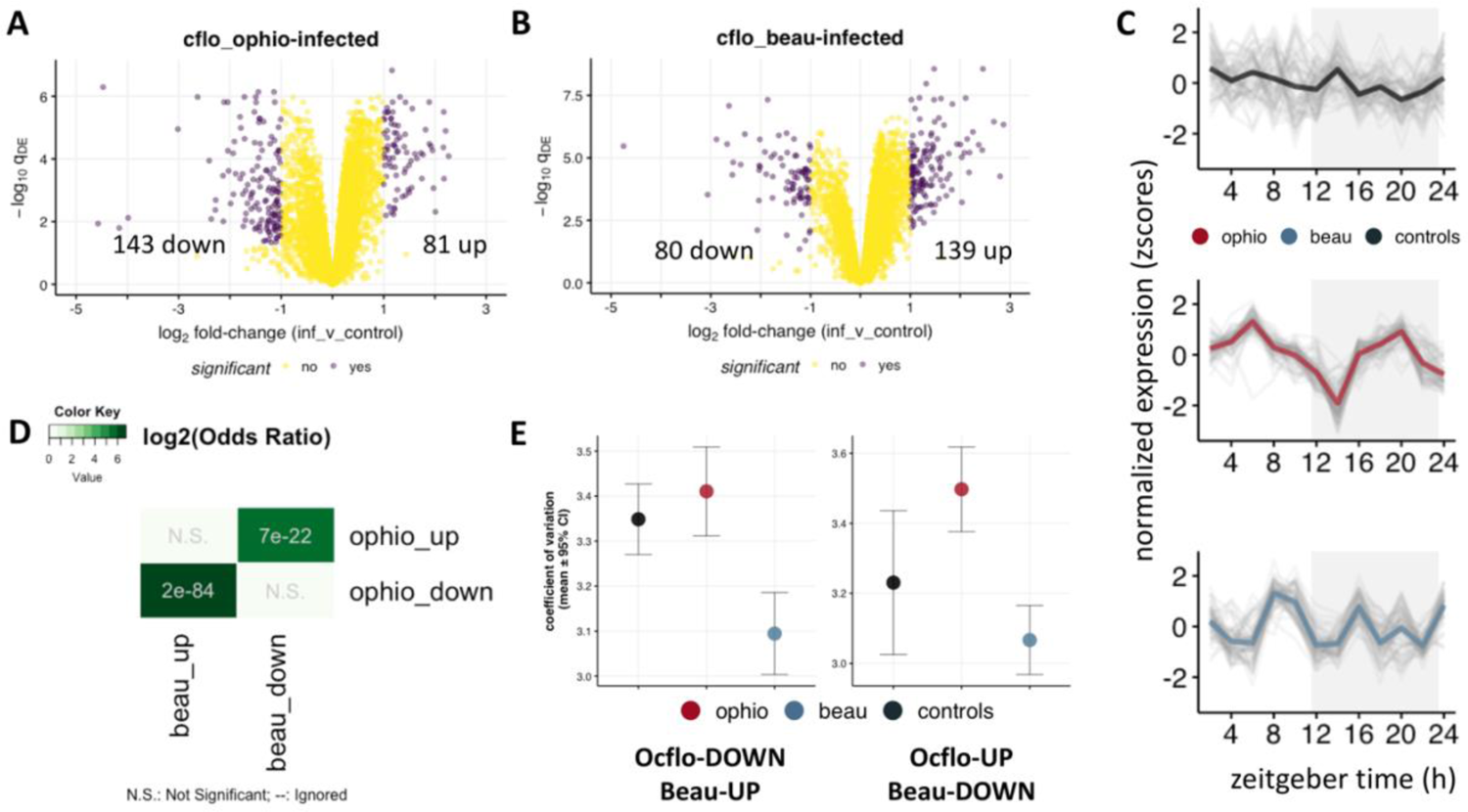
Differential gene expression reveals two core sets of ant genes that are affected during fungal infections, but the direction of the effect is parasite dependent. The volcano plots show the results of differential gene expression analysis for (A) O. camponoti-floridani-infected ant heads and (B) B. bassiana-infected ant heads, as compared to controls. Purple dots indicate genes that were classified as significantly differentially expressed (fold-change ≥ 2, p-value < 0.05). The number of up and down regulated genes during an infectious disease, as compared to controls, are shown. Panel (C) shows the stacked zplots for the 63 genes that were found to be significantly down regulated in O. camponoti-floridani-infected ants but up regulated in ant heads during B. bassiana infections (Ocflo-Down, Beau-UP). Each grey line represents daily expression of one gene, and the colored solid line represents the median daily expression for all genes in the set. The x- and y-axis of the stacked zplot have the same meaning as before. Panel (D) shows the results of pairwise Fisher’s exact test. The color of each cell indicates the log2-(odds-ratio) for the overlap between two gene sets and the p-values used to infer significance of overlap are shown. The odds ratio and p-value are only shown for significant overlaps (p-value < 0.05). Panel (E) shows the amplitude of the genes found to be (left) significantly down regulated in O. camponoti-floridani-infected ants but up regulated in B. bassiana-infected ants (Ocflo-DOWN, Beau-UP), and (right) significantly up regulated in O. camponoti-floridani-infected ants but down regulated in B. bassiana-infected ants (Ocflo-UP, Beau-DOWN), as compared to controls, in all three treatment groups. The circles show mean amplitude of the respective genes and the error bars indicate 95% CI.

Looking at the identity of these 63 *Ocflo*-DOWN but Beau-UP genes (downregulated during *O. camponoti-floridani* infection but upregulated during *B. bassiana* infection), we found several genes-of-interest: *bubblegum* (affects neurodegeneration and lifespan in flies; [53, 54]), pheromone-binding protein *Gp-9-like* (regulator of social complexity in ants; [55]), *D-3-phosphoglycerate dehydrogenase* (caste-associated DRG in *C. floridanus*; [1]), venom acid phosphatase *Acph-1-like* (a known venom acid phosphatase; [56]), the juvenile hormone esterase *venom carboxylesterase-6* (juvenile hormone esterase that show caste-associated DRG in *C. floridanus*; [1]), three copies of *juvenile hormone acid O-methyltransferase* (catalyzes the last step of JH synthesis to regulate titers of both JH and Vg in insects; [57]). This relatively small set of genes showed significant overrepresentation for *cytochrome p450* genes involved in oxidoreductase activity, iron ion binding, and heme binding.

Furthermore, the *Ocflo*-DOWN-Beau-UP genes showed synchronized daily expression in *O. camponoti-floridani*-infected ants with a sharp dip at ZT14 and seemingly bimodal peaks (Fig. 4C). In fact, most of the genes downregulated in *O. camponoti-floridani*-infected ants showed a similar synchronized rhythmicity during *O. camponoti-floridani* infection with a bimodal peak of daily expression, suggestive of 12h oscillations. In comparison, for *Ocflo*-UP genes, relatively less synchronization of their daily fluctuations was found, although most *Ocflo*-UP genes showed a clear local trough at ZT6. The presence of 12h oscillations in *Ocflo*-DOWN genes during *O. camponoti-floridani* infections was confirmed as we found a significant overlap of *Ocflo*-DOWN genes with ones classified as 12h-rhythmic in *O. camponoti-floridani*-infected ants. Of more interest is the timing of the synchronized sharp dip observed around ZT14 for these *Ocflo*-DOWN genes (Fig. 4C, Supp. 3B), the same time of day at which majority of the rhythmic genes and processes in *O. camponoti-floridani*-infected ants show a peak activity (Fig. 3B). It seems, therefore, that at the halfway mark in *O. camponoti-floridani* infections, host genes are significantly downregulated as compared to controls, in a time-of-day specific manner, and the timing of this downregulation correlates with the peak time of activity for most genes found oscillating in the infected ant head.

Taken together, our findings suggest a possible mechanism via which *O. camponoti-floridani* might be regulating the phase of the host’s clock-controlled genes, and the processes they regulate. *O. camponoti-floridani* is known to secrete toxins, including several enterotoxins, while inside their ant host, some of which are rhythmically expressed [16]. We do not have the data to test if during infection, the timing of peak toxin activity, driven by the parasite’s clock, correlates with the time at which we observe a sharp deactivation of host genes involved in oxidoreductase activity. However, if we do find that the correlation exists, it would be suggestive that at least some of these mycotoxins likely function to reduce the oxidative stress in the ant host but in a timely manner. Given that the cluster of downregulated genes we have identified in *O. camponoti-floridani*-infected ants contains several behavior regulatory genes (juvenile hormone esterases), the parasite’s ability to manipulate host behavior in a timely manner might have evolved due to pre-existing links between host’s behavioral state and response to oxidative stress.

In comparison, the relatively fast progression of *B. bassiana* infections in ants might be, in part, due to a significantly upregulated host response to oxidative stress. Future studies comparing the changes that occur to the parasites’ daily transcriptome during disease, and how the phase of rhythmic fungal effectors compare with the host’s response to oxidative stress will be necessary to shed light on our speculations. However, for our argument to hold, the timely changes in the DEGs would need to have time-of-day effects on the host’s clock-controlled processes. In other words, there needs to be cross-talk between the host’s behavior regulatory genes and clock output, and that’s exactly what we have previously found [1].

### Effects of infectious diseases on host gene expression network

Thus far, we have compared our time-course transcriptomics datasets to identify several sets of genes that characterize the disease-associated changes in the host’s daily transcriptome. In this section, we take a systems approach to identify the regulatory mechanisms that potentially link these genes of interest. Furthermore, constructing the global network of gene expression in uninfected controls, and characterizing the changes to the network during disease allowed us to ask, for example, if the accelerated disease progression of *B. bassiana*, a necrotrophic parasite, is due to substantial changes to the structure of the host’s expression network. In comparison, infection by *O. camponoti-floridani* might cause less drastic but targeted changes to certain parts of the network, with the affected regions potentially being overrepresented for core clock and clock-controlled genes. Most importantly, using network analyses we aimed to integrate all of our findings, from the current study as well as prior studies done in our lab, in order to put forward a hypothesis for the regulatory mechanism via which manipulating parasites such as *O. camponoti-floridani* might be driving time-of-day specific changes in their host’s behavior.

Prior to this study, we used meta-analysis to show that genes differentially expressed between foragers and nurses (behavioral plasticity genes; [1]) were primary located in two modules of the ant’s brain gene expression network [4]. Additionally, we discovered that both behavior plasticity modules were connected to at least one rhythmic module of the host. This suggested that in ants, at least in *C. floridanus*, there exists a putative regulatory link between genes underlying behavioral plasticity (caste-associated DEGs) and clock-controlled rhythmic output. Therefore, drastic changes to the expression of these two behavioral plasticity modules, due to changes in the ant’s social context or disease state, could induce changes to the rhythmic modules and the different processes they regulate, including rhythmic behavior.

In a separate study, we had identified over a thousand genes that were differentially expressed in ant heads at the manipulated biting stage (infected ants biting down on a substrate with a locked jaw) prior to death. When we mapped the genes differentially expressed in the ant heads during active manipulation (biting) onto the ant’s brain gene expression network, we found that most of these manipulation genes were located in exactly the same modules as the behavioral plasticity genes.

This led us to hypothesize that, to induce timely behavioral changes in its host, *O. camponoti-floridani* likely targets existing molecular links between host’s behavioral and chronobiological plasticity. Here, we empirically test this hypothesis using the time-course RNASeq data that we have collected for infected ant heads at the halfway mark in both diseases. For our hypothesis to hold, we should observe the following in our data, (1) the presence of molecular links between behavioral plasticity and rhythmic genes (i.e., for-UP or for-DOWN modules connected to rhythmic modules in the gene expression network), and (2) *O. camponoti-floridani* infections significantly affecting the expression of the behavioral plasticity genes (i.e., *Ocflo*-UP (DOWN) genes and for-UP (DOWN) genes should be located in the same modules).

We constructed and annotated the gene co-expression network (GCN) of control forager heads as per the protocol detailed in de Bekker and Das (2022) [4]. We built the ant’s GCN using the 9591 genes that were expressed (≥1 FPKM) in control ant heads for at least half of all sampled time points. We wanted to use the gene expression network of uninfected controls as the background to identify the changes that occur in the host’s network during infection. But building the ant’s network using only genes that are expressed in control conditions did not allow us to map the genes that might be expressed during infection, but not in controls, on to this network, which is a limitation of our analyses. However, the reference network we built should allow us to achieve the goal of identifying the modules that show an overlap with our genes of interest. These 9591 expressed genes were clustered into 22 modules based on co-expression and named accordingly (C1 to C22). The global connectivity patterns of the ant GCN is shown in Figure 5A, where nodes represent the modules or clusters of highly co-expressed genes (Kendall’s tau-b correlation ≥ 0.6), and edges between modules indicate a similarity of module-module expression (Kendall’s tau-b correlation of at least 0.6 between a module’s eigengene expression with another).

**Figure 5.**
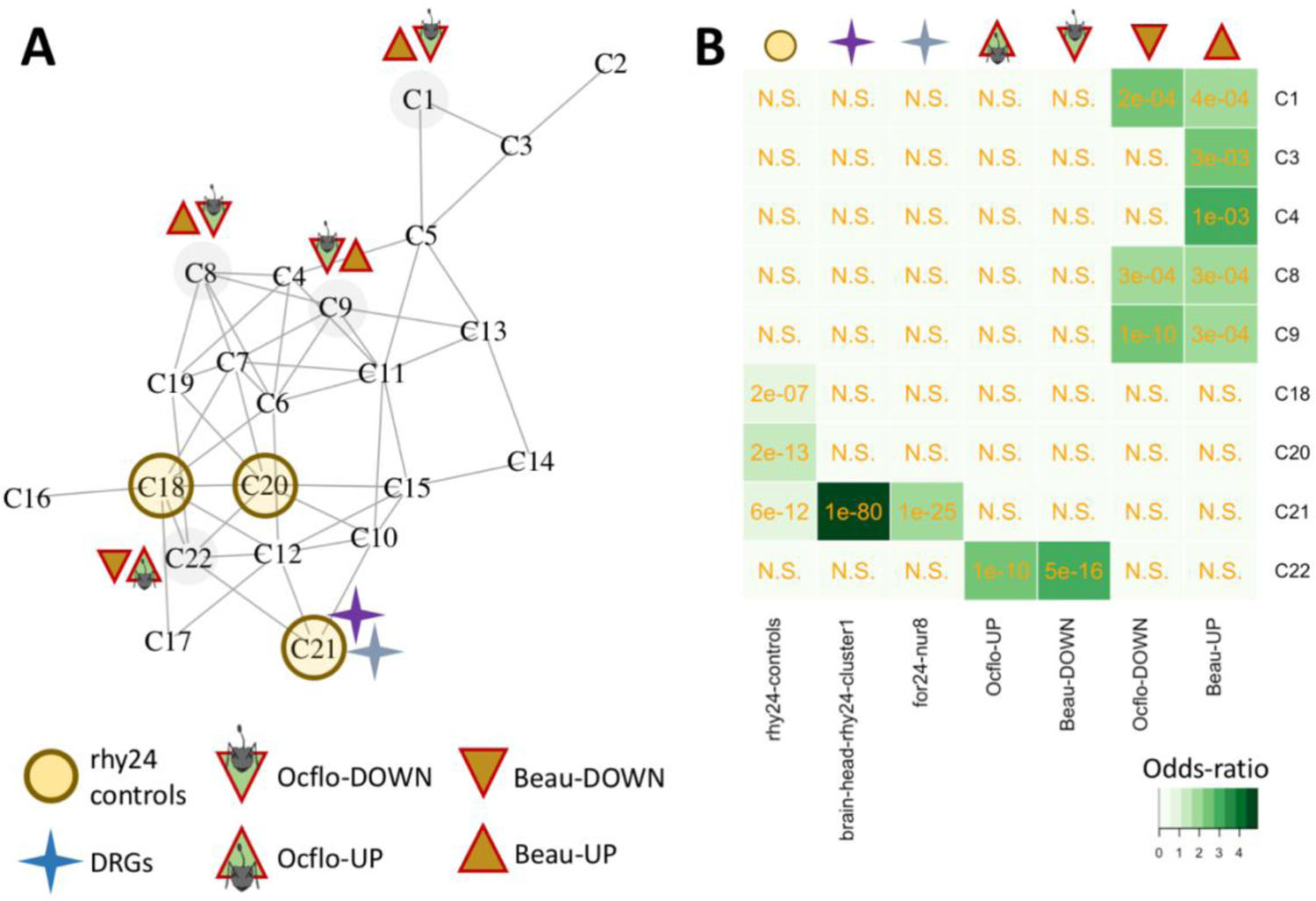
Annotated gene co-expression network of uninfected forager heads. Panel (A) shows the annotated gene co-expression network (GCN) of uninfected (control) forager heads of *Camponotus floridanus*. The annotations in the network shows the different modules of interest that we have identified as being putatively important for the cross-talk between ant’s clock-controlled rhythmicity, behavioral plasticity, and disease-associated differential expression and differential rhythmicity. The nodes, labelled C1 through C22, represent modules of highly co-expressed ant genes, and the edges represent co-expression of connected modules. All edges indicate module-module correlations between 0.6 and 0.8, and no edges between two modules indicate correlations <0.6. The abbreviations have the following meaning: (i) rhy24 controls indicate modules that show an overrepresentation for ant genes that were classified as significantly rhythmic in uninfected controls, (ii) DRGs indicate the modules that show overrepresentation for ant genes that show disease-associated (purple) and caste-associated (light blue) differential rhythmicity (brain-head-rhy24-cluster1 and for24-nur8, respectively), (iii) *Ocflo*-DOWN (or *Ocflo*-UP) identifies the module(s) in the ant’s GCN that shows a significant overrepresentation for genes down regulated (or up regulated) in ant heads at halfway through *O. camponoti-floridani* infections, and (iv) Beau-DOWN (or Beau-UP) identifies the module(s) in the ant’s GCN that shows a significant overrepresentation for genes down regulated (or up regulated) in ant heads at halfway through *B. bassiana* infections. Panel (B) shows the results of the pairwise Fisher’s exact test that was performed to identify the modules of interest in (A). The respective gene sets have been discussed in detail within the main text.

**Figure 6.**
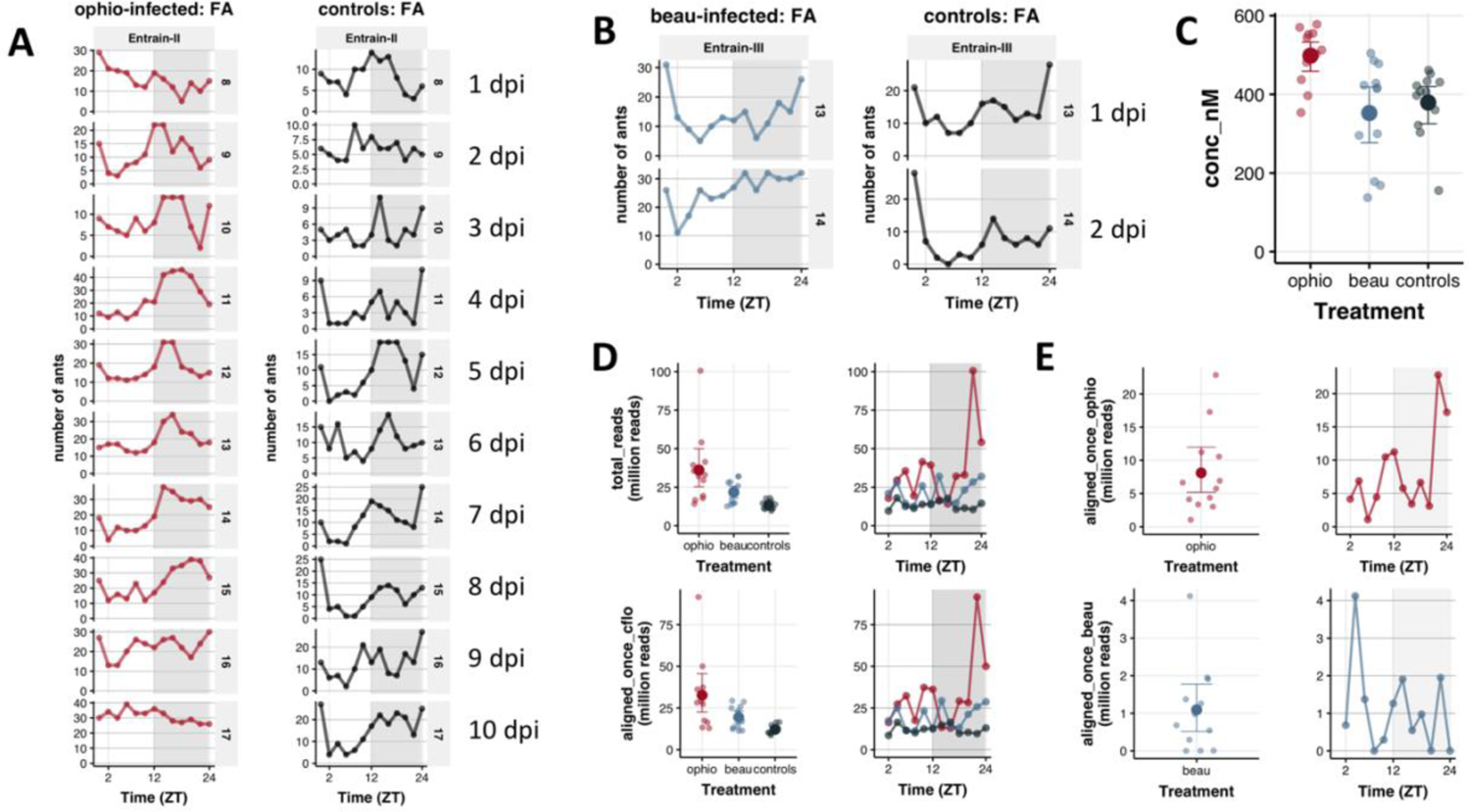
Ant activity rhythms and quality of time-course transcriptomes. The daily foraging activity of ants infected with (A) *Ophiocordyceps camponoti-floridani* (red) and (B) *Beauveria bassiana* (blue) during the entrainment period post infections and prior to sampling are shown. For each infection run, the daily foraging activity of the uninfected controls that were monitored in parallel are also shown (black lines). Once sampled, total RNA was extracted from the whole heads of infected ants and uninfected controls for RNASeq. For each treatment group, (C) shows the concentration (in nM) of extracted total RNA from each sample (lighter circles), along with the mean (darker circles) and 95% confidence intervals (CI) for each treatment group. (D) shows the total number of reads (in million) obtained for each sample, along with their mean and 95% CI (top row), as well as the number of reads that uniquely mapped to the *C. floridanus* genome (bottom row). Given that sequencing the infected ant heads resulted in mixed transcriptomes containing both ant and fungal reads, (E) shows the number of reads from the respective mixed transcriptomes that mapped to the *O. camponoti-floridani* and *B. bassiana* genome.

Next, we annotated the GCN of forager heads by identifying where in the network our genes of interest are located to understand how these genes regulate each other. In addition to testing our aforementioned hypothesis, annotating the GCN also allowed us to ask, for example, if drastic changes in the expression of a set of genes (DEGs) during disease might be able to induce differential rhythmicity (synchronized changes in the daily expression) of core clock components, clock modulators, or clock-controlled output. To annotate the ant GCN, we identified the module(s) that have a significant overlap with genes displaying (1) significant 24h rhythms in control heads (rhy24h-controls), (2) disease-associated differential rhythmicity (brain-head-rhy24-cluster1, Fig. 2B), (3) caste-associated differential rhythmicity (for24-nur8, discussed in *Changes in ant daily gene expression during infection: Differentially Rhythmic Genes*; from [1]), (4) significant differential expression between control and diseased states (*Ocflo*-UP/DOWN and Beau-UP/DOWN; Fig. 4A-B), and (5) significant differential expression between foragers and nurses (for-UP/DOWN, also referred to as behavioral plasticity genes; from [1]).

We found that most of the genes identified as 24h-rhythmic in controls were primarily located in three modules (C18, C20, and C21) (Fig. 5B). To validate our annotation, we looked at the identity of the genes clustered into each of the rhythmic modules as we expected to see known insect clock components in these three clusters. Indeed, we found that module C21 contained core clock genes (*Clock* and *Period*), clock modulators (*Doubletime* and *Nemo*), and clock output genes (*mAchR, DopEcR, Pka*, and *Lark*) (Fig. 15). Additionally, C19 contained the clock modulating kinases *Ck2a* and *Pp1*, whereas C18 contained the gene encoding the major insect photoceptor *Rhodopsin* (Additional File 1).

We should note, not all of the aforementioned genes were classified as significantly 24h-rhythmic by eJTK. We believe this inconsistency is likely due to the choice of an arbitrary p-value cutoff to classify genes into rhythmic and otherwise, and not the absence of rhythmicity. We briefly explain our reasoning. A network-based identification of rhythmic modules is a powerful tool as it allows us to identify clusters of rhythmic genes using guilt-by-association. As we have discussed previously, the modules in the ant GCN were created on the basis of co-expression, i.e., how similarly do two genes mirror each other’s daily expression pattern. Therefore, if a significant number of genes in a module are classified as rhythmic, then the probability that majority of the genes in the module (cluster of highly co-expressed genes) are rhythmic is relatively high. The presence of known clock components in rhythmic modules, therefore, likely suggest that these clock genes are also rhythmic in uninfected controls, consistent with our expectations.

Having identified the rhythmic modules in our ant GCN, we wanted to know if the structure (gene-gene connectivity patterns) of the rhythmic modules changed drastically during disease or not. For a given infectious disease, we calculated each module’s preservation using the Preservation Zsummary score proposed by Langfelder and colleagues [42]. As the author’s point out using simulations, a Zsummary score of greater than 10 indicates that there is strong evidence that the structure of the module is preserved, a Zsummary score between 2 and 10 indicates moderate evidence for module preservation, whereas Zsummary < 2 provides no evidence for module preservation. To be stringent, we classified a module to be preserved if it showed Zsummary > 10, but not preserved otherwise. Comparing the ant transcriptome during *B. bassiana* infections to controls, we found only three modules that showed a Zsummary > 10, all of which were rhythmic modules (preservation ranking: C21 > C20 > C19). In *O. camponoti-floridani*-infected ants, however, only two modules showed evidence for preservation, one of which was the rhythmic module C21, also found to be preserved during *B. bassiana* infections.

Given that C21 contains core clock genes as well as clock modulators, our finding hints at the possibility that during either disease the connectivity patterns of the core feedback loop of the host clock does not get drastically altered. However, ants infected with *O. camponoti-floridani* do seem to lose their daily foraging rhythms at halfway through disease progression, but *B. bassiana*-infected ants do not. This might be due to the fact that we find that all three of the host’s rhythmic modules show a high degree of preservation during *B. bassiana* infections, at least in their connectivity patterns. However, it does not eliminate the possibility of an altered or differential co-expression, the occurrence of which would explain the altered pattern of clock-controlled foraging rhythm that Trinh and colleagues have observed in ants during late-stage *B. bassiana* infections. The loss of locomotory rhythms in *O. camponoti-floridani*-infected ants, therefore, might be due to drastic rewiring that occurs in either one or both of the rhythmic modules (C18 or C20) which contains the photoreceptor *Rhodopsin* and the kinases *Ck2a* and *Pp1*. We should point out the *O. camponoti-floridani* clock gene *CK2*, which functions as a core clock gene and a photoreceptor in the model fungi *Neurospora*, shows a significant upregulation throughout the disease progression, both at the halfway mark as well as the manipulated biting stage (discussed in detail in [16]). If this finding suggests a direct interaction of the fungal-ant clocks during disease or not remains unclear, but the possibility remains. In comparison, at halfway through both diseases, the rhythmic module C21 containing core clock genes *Per* and *Clk* show a high degree of preservation, highlighting that host genes that tightly co-express with its core clock components are likely essential for host survival, and as such remain unaltered as the parasite incubates inside its host.

Most of the ant genes significantly up-regulated during *O. camponoti-floridani*-infection were located in one module only (C22), and the same module contained most of the genes down-regulated during *B. bassiana*-infection. In comparison, ant genes significantly downregulated during *O. camponoti-floridani* infections were overrepresented in three modules (C1, C8, and C9), all of which also showed overrepresentation for genes upregulated in *B. bassiana*-infected ants. This reconfirms our conclusion about the DEGs as discussed in the previous section. There seems to exist a core set of genes that are differentially expressed during both diseases (disease associated DEGs). What distinguishes one infectious disease from another is the direction of this differential expression. As we have discussed above, our prior meta-analyses revealed that *O. camponoti-floridani* might be inducing differential expression of the same set of ant genes that underlie behavioral plasticity (DEGs between foragers and nurses).

Mapping the behavioral plasticity genes on to our ant GCN, we found that the genes downregulated in foragers (versus nurses; for-DOWN) were mostly located in module C1 whereas genes upregulated in foragers (for-UP) were located primarily in module C22, the same modules that show overrepresentation for disease-associated DEGs. Therefore, it appears that expression levels of C1 and C22 genes are associated with behavioral plasticity as well as parasite-specific disease outcomes. More specifically, genes in module C22, usually upregulated in foragers (versus nurses), show an even stronger upregulation during *O. camponoti-floridani* infection. Whereas, during *B. bassiana* infection, the same cluster of genes show a significant downregulation. To understand what function such upregulation of C22 (or downregulation of C1) might be serving in *O. camponoti-floridani*-infected ants, we first need to discuss the possible links between differential gene expression and differential rhythmicity as evidenced by our annotated GCN.

We found that module C22, that contains the majority of *Ocflo*-DOWN, *Beau*-UP, and for-UP genes, is connected to all three of the rhythmic modules (C18, C20, C21) in our ant GCN (Fig. 15). However, we did not find any edges connecting module C1 to any of the rhythmic modules. Therefore, it seems that module C22 might be functioning as a regulatory switch in ants that modulates the rhythmic properties of the clock and its output. When we mapped, respectively, the genes that show caste-associated and disease-associated differential rhythmicity on to the GCN, we found that both sets of genes showed a significant overlap with only one module: C21. Therefore, it appears that drastic changes (up/down-regulation) in the expression of C22 could induce differential rhythmicity of C21 genes which contains essential clock genes and modulators.

In summary, the comparative network analyses have shown that (1) molecular links between behavioral plasticity and rhythmic genes that we found for ant brains also exist at the scale of whole heads, (2) not only *O. camponoti-floridani*, but also *B. bassiana*, significantly affects the expression of modules overrepresented for behavioral plasticity genes, (3) the direction of the effect (up or downregulation) on these behavioral plasticity genes is exactly the opposite for infections by *O. camponoti-floridani* and *B. bassiana*, and (4) disease associated changes to the expression of host genes regulating behavior plasticity, on the forager-nurse axis, is correlated with synchronized changes in the daily expression of clock genes including *Period, Clock, Ck2a*, and *Pp1b*.

## CONCLUSION

We started our investigation with the apriori hypothesis that *O. camponoti-floridani* affects the biological clock and clock-controlled rhythms of infected ant host. Given that changes in host’s clock-controlled rhythms might be a general hallmark of infectious diseases, we compared the changes in daily transcriptome of *C. floridanus* ants while infected with the manipulating parasite *O. camponoti-floridani* versus a generalist, non-manipulating fungi *Beauveria bassiana*. Using time-course transcriptomics and comparative network analyses, we found that changes to the host’s clock-controlled gene expression might be a general hallmark of infectious disease as both fungal parasites induced differential rhythmicity in the same set of rhythmic host genes. The study puts forward several data-driven hypotheses for the regulatory mechanisms via which *O. camponoti-floridani* might be inducing changes to the host’s clock and clock output, including clock-controlled behavior. The consistent regulatory patterns and genes of interest that we have identified in this study can serve as a repository of candidate host targets that might play a role in modifying host behavior, at least for *Ophiocordyceps*-ant systems.

## SUPPLEMENTARY FILES

### Additional File 1

#### Results of the network analyses

The following CSV file contains the results from the network analysis and includes the list of all ant genes used to build the gene co-expression network, their module identity, functional annotations, and all relevant information that is necessary to follow the main text.

#### Link to file

https://github.com/biplabendu/Das_et_al_2022a/blob/master/manuscript/99_dissertation/07_Ocflo_GCN_results.csv

### Additional File 2

#### Validation of sampling time point at halfway through disease progression

In this study, we aimed to look at the daily rhythms in gene expression of infected ants halfway through disease progression since this is where the daily foraging activity of carpenter ants infected with *Ophiocordyceps* and *B. bassiana* appears to be disturbed, albeit in a different manner [15]. We validated our sampling timepoint by comparing daily foraging activity of the infected ants in each of our treatment groups (*O. camponoti-floridani*-infected and *B. bassiana-* infected ants) to that of Trinh and colleagues [15]. We did so by counting the number of ants present in the 12h:12h LD exposed foraging arena, every 2h, throughout the experiment (after initial infections and prior to sampling). The entrainment cues (temperature, humidity, and light levels) presented in the foraging arena were consistent across the two infection runs (Supp. Fig. 1). Therefore, any time-of-day specific differences that we found in *O. camponoti-floridani*-infected as compared to *B. bassiana*-infected ants are likely to be the result of infection differences and not due to differences in their external environment in terms of temperature, humidity, or light. The uninfected controls were exposed to entrainment cues identical to the ones shown for the two infection runs.

**Supplementary Figure 1.**
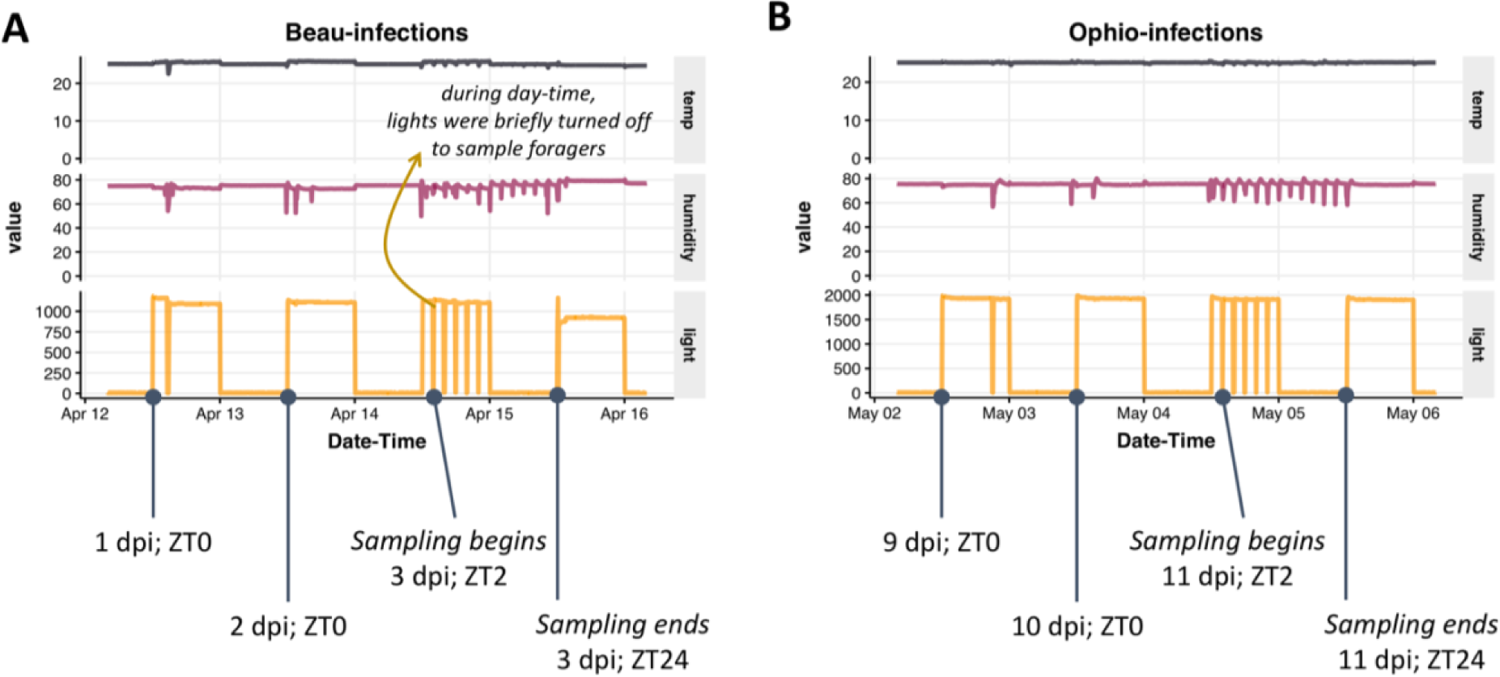
Environmental conditions in the foraging arena during infection run. The abiotic conditions (light intensity, temperature, and relative humidity) in the foraging arena of the experimental setup are shown for ants infected with (A) *Beauveria bassiana* (Beau) and (B) *Ophiocordyceps camponoti-floridani* (Ophio). Data is shown for two days prior to sampling, the sampling day, and one day after sampling.

We quantified the daily foraging activity of the *O. camponoti-floridani*-infected and *B. bassiana*-infected ants in parallel to uninfected controls. As shown in Supplementary Figure 2A, *O. camponoti-floridani*-infected ants maintained daily foraging rhythms until 6 to 7 days post infection (dpi), and by 10 dpi their foraging levels showed no apparent daily rhythm. This loss of rhythmicity coincided with the halfway point of infection during which only half of all ants infected with *O. camponoti-floridani* survived (i.e., lethal time (LT) 50%), which is consistent with our prior knowledge [15]. As such, we sampled *O. camponoti-floridani*-infected ants on 11 dpi. Corresponding samples of *B. bassiana*-infected ants at halfway through disease were sampled on 3 dpi. Uninfected control ants were sampled in conjunction with *B. bassiana*-infected ants. Both their daily foraging activities prior to sampling are shown in Supplementary Figure 2B.

### Additional File 3

#### Quality of transcriptomes and possible sources of variation

To obtain RNA for sequencing, we pooled three technical replicates (whole heads), per treatment group, for each time point of sampling. What we aimed to obtain in the process is the average gene expression for a given treatment group, for each time point. Given that we were working with only one value per time point, per treatment, we wanted to ensure that time-of-day specific patterns that we find in our analyses were not introduced by differences in sample processing and sequencing. Therefore, we checked the concentration of cDNA libraries used, number of reads sequenced per sample, and number (and percent) of reads mapping to the *Cflo* genome.

The concentrations of all cDNA libraries were comparable (controls = 379 ± 25 nM, *O. camponoti-floridani*-infected ants = 498 ± 20 nM, and *B. bassiana -*infected ants = 352 ± 38 nM; mean ± standard errors) (Supp. Fig. 2C). Performing RNASeq, we aimed to produce more reads for infected individuals than for controls since we needed to account for the portion of fungal reads in the mixed transcriptomes that we would obtain [11]. As such, we obtained 13.4 (± 0.8) million reads for control ants, 36.1 (± 0.7) million reads for *O. camponoti-floridani*-infected ants, and 21.8 (± 0.2) million reads for *B. bassiana-*infected ants. On an average, 12.2 (± 0.8) million reads for control ants, 32.7 (± 6.2) million reads for *O. camponoti-floridani*-infected ants, and 19.5 (± 2) million reads for *B. bassiana*-infected ants mapped uniquely to the ant genome (Supp. Fig. 2D). This is consistent with the advised coverage depth to reliably detect daily rhythms in gene expression [58]. As such, our environmental conditions, sampling time point and sample processing for RNASeq should have resulted in a dataset that would allow us to investigate infection-related differences in daily gene expression in *C. floridanus* ants.

### Additional File 4

#### Fungal gene expression inside ant heads

We found substantial amounts of *O. camponoti-floridani* mRNA in the ant heads at halfway through disease progression (8.1 ± 1.8 million reads from the mixed transcriptome mapped uniquely to the *O. camponoti-floridani* genome) (Supp. Fig. 2E). Whereas for *B. bassiana*-infected ants, we did not find much fungal mRNA in ant heads at the same disease stage (only 1.1 ± 0.3 million reads mapped uniquely to *B. bassiana* genome). This difference in fungal load inside ant heads might be reflective of the different strategies employed by the two fungal parasites to ensure successful infection in its ant host. *B. bassiana* is a generalist necrotrophic parasite that kills its ant host within in a week, and as such might not require entry into the ant’s head for ensuring infection and eventual transmission. In comparison, *O. camponoti-floridani* behaves as a hemibiotroph, growing inside its alive host for weeks and requiring timely manipulation of the host’s behavior for successful transmission (reviewed in [14]). It has been shown previously that *Ophiocordyceps* do grow into its host’s head and mandibular tissues [9], and the proximity of fungal growth inside the head, around the ant brain, has been hypothesized to be important for successful manipulation of ant behavior [59]. We further investigated *O. camponoti-floridani* daily gene expression during infection and how it compares to the daily expression exhibited in liquid culture outside the host (discussed in [16]).

## Notes

### Competing Interest Statement

The authors have declared no competing interest.

### Summary of Updates

Author ORCIDs have been added. No change has been made to the manuscript.

